# Grid cell co-activity patterns during sleep reflect spatial overlap of grid fields during active behaviors

**DOI:** 10.1101/198671

**Authors:** Sean G. Trettel, John B. Trimper, Ernie Hwaun, Ila R. Fiete, Laura Lee Colgin

## Abstract

Continuous attractor network models of grid formation posit that recurrent connectivity between grid cells controls their patterns of co-activation. Grid cells from a common module exhibit stable offsets in their periodic spatial tuning curves across environments, which may reflect recurrent connectivity or correlated sensory inputs. Here we explore whether cell-cell relationships predicted by attractor models persist during sleep states in which spatially informative sensory inputs are absent. We recorded ensembles of grid cells in superficial layers of medial entorhinal cortex during active exploratory behaviors and overnight sleep. Per pair and collectively, we found preserved patterns of spike-time correlations across waking, REM, and non-REM sleep, which reflected the spatial tuning offsets between these cells during active exploration. The preservation of cell-cell relationships across states was not explained by theta oscillations or CA1 activity. These results suggest that recurrent connectivity within the grid cell network drives grid cell activity across behavioral states.

---

Grid cells of the medial entorhinal cortex (MEC)^1^ together with place^2^, head direction^3,4^, border^5^, speed cells^6^, and cells that simultaneously code multiple navigational variables^7^, convey information about the evolving location and orientation of mammals as they move through 2D open fields, run on 1D linear tracks, or fly through 3D space^8^. Grid cells are defined by regular, periodic responses to an animal’s 2D spatial location. Each grid cell’s multiple spatial receptive fields (“grid fields”) form a characteristic geometric pattern well-described by a lattice of equilateral triangles.

Several different classes of models seek to explain grid cell activity and function^9,10,11,12,13,14^. One of the principal models is based on continuous attractor dynamics that emerge through pattern formation in networks with strong lateral connectivity^9,11,15^. In such continuous attractor network models, connectivity between cells tightly constrains their patterns of co-activation, and thus connectivity is closely related to relationships in spatial tuning. Specifically, these models predict that cell pairs with strong short-time correlations should exhibit similar spatial tuning phases, whereas most pairs with weak short-time correlations or negative correlations should exhibit offset or anti-phase spatial tuning. Moreover, these correlations should be preserved across states because the circuit dynamics are always shaped by the same recurrent connectivity in the network. This prediction can be tested in recordings of the same grid cell ensembles across active waking, quiescent, and sleep states.

Other spatially responsive cells appear to have structured activity patterns during sleep. During waking rest and non-REM sleep (NREM), place cells show a transient increase in their spike time correlations that is related to the firing patterns of the place cells during exploration^16,17,18,19^. Head direction cells have been shown to exhibit structured activity during sleep states as well^20,21^. Considering that place cells, head direction cells, and grid cells are part of a larger navigational and spatial memory system, and that grid cells in the superficial layers of the MEC provide input to place cells^22,23,24,25^, it is possible that grid cells in the superficial layers of MEC drive hippocampal place cell reactivation during sleep. However, little is known about superficial layer MEC grid cell firing patterns across overnight sleep, and a recent study suggested that coordinated reactivation typically does not occur between place cells and grid cells in the superficial layers of MEC^26^.

The central question of this paper is to ask whether, and to what extent, states in the grid cell circuit across sleep resemble those during waking. These findings will help to better understand the potential role of the grid cell circuit in driving dynamics of the larger navigational and spatial memory system. Grid cells are hypothesized to integrate velocity cues as an animal navigates through an environment, thus obtaining a continuously updated estimate of location even when spatially informative external cues are unavailable^1,27,28^. Also, grid cells have more recently been shown to play a role in non-spatial tasks, namely measuring the passage of time^29^ or creating a map of the visual field^30^. Although grid cells have been characterized as exhibiting grid-like responses during both spatial and non-spatial tasks, the presence or stability of relationships between grid cell pairs across both spatial and non-spatial tasks has not been examined. Thus, it remains unclear whether the cell-cell relationships predicted by the continuous attractor network models persist across spatial and non-spatial behaviors.

Here, we examined spike time cross-correlations in large numbers of MEC grid cell pairs in freely behaving rats during exploratory behaviors and across several hours of overnight sleep. Hippocampal circuits have been shown to exhibit preserved patterns of very recent past experience during the first 20 mins of sleep^17,19^. In contrast, we examine a prediction of low-dimensional continuous attractor models, namely that patterns of activity during sleep in grid cells exhibit a preserved structure across hours of overnight sleep. We found that spike-time and spike-rate correlations between grid cells were related to the degree of overlap in their spatial tuning curves and that, remarkably, these relationships persisted during rapid eye movement (REM) and nonREM (NREM) sleep. Moreover, the preserved patterns of co-activity during sleep were not explained by theta entrainment of spike times or analogous hippocampal co-activity patterns.

## RESULTS

### Activity of grid cells during waking and sleep

To examine the structure of the grid cell circuit responses during sleep, we simultaneously recorded multiple single units in MEC over several hours as rats (n = 6) ran in open field and across the entirety of their overnight sleep cycle. From these ensembles, we identified putative grid cells using a gridness score^3^ (see Methods) calculated from three twenty-minute open field sessions. Grid cell recordings were only obtained from MEC superficial layers (i.e., II and III). Grid cells were only included if they remained stable across active waking recordings, subsequent overnight sleep recordings, and additional open field recordings the next morning (see Methods). From a total of 157 putative grid cells and 417 possible putative grid cell pairs, 226 grid cell pairs from 6 animals passed our inclusion criteria (see Methods; Table 1).

**Table 1.** Grid cell counts per animal.

| **<u>Animal</u>** | **<u>Number of Grid Cells</u>** | **<u>Number of Grid Cell Pairs Used</u>** | **<u>Number of Recording Days</u>** |
| --- | --- | --- | --- |
| **Rat 20** | 33 | 57 | 5 |
| **Rat 26** | 24 | 52 | 4 |
| **Rat 29** | 5 | 9 | 1 |
| **Rat 74** | 20 | 10 | 7 |
| **Rat 78** | 24 | 78 | 5 |
| **Rat 109** | 52 | 75 | 7 |

Grid cells were active during spatial exploration of open field environments and during both REM and NREM stages of sleep (Figure 1). However, firing rates were significantly different across the three behavioral states (157 grid cells; RUN: 2.04 +/− 0.10 Hz, REM: 1.08 +/− 0.06 Hz, NREM: 0.76 +/− 0.04 Hz, F(2, 468) = 89.6, p < 0.0001; 1-way ANOVA; Figure 1b-c). Firing rates during REM and NREM were significantly lower than during waking behaviors (p < 0.0001 in both cases, post hoc Tukey’s HSD test). In addition, REM firing rates were significantly higher than NREM firing rates (p = 0.004, post hoc Tukey’s HSD test).

**Figure 1.**
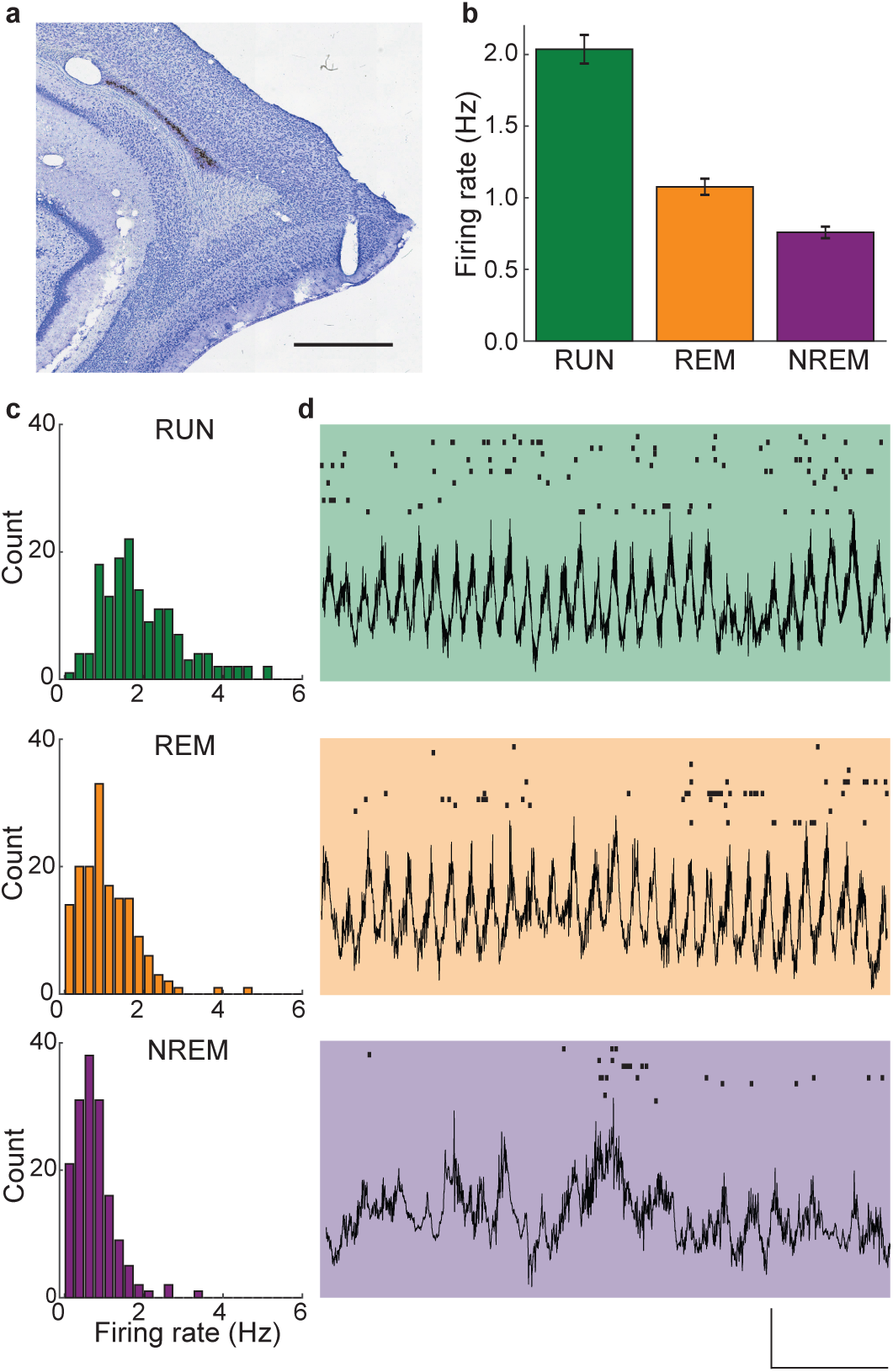
Grid cell firing rates varied across active and sleep states. (a) An example histological section shows a tetrode track in the superficial layers of the medial entorhinal cortex. Scale bar is 1 mm. (b) Grid cell firing rates differed significantly across behavioral states. Firing rates (mean +/− SEM) were highest during active ambulation and lowest during NREM sleep. (c) Histograms show the distributions of firing rates across grid cells segregated by behavioral state. (d) Representative examples of LFP epochs are presented with simultaneous spike rasters from an example ensemble of grid cells plotted above each trace. Each row of the raster plots corresponds to a single grid cell. Activity from the same ensemble of grid cells is shown in each behavioral state. Vertical scale bar is 500 μv, horizontal scale bar is 1 s.

### Patterns of co-activation in individual grid cell pairs were maintained across active and sleep states

We next investigated whether co-activity patterns of pairs of grid cells were related to overlap in their spatial tuning properties, as predicted by attractor network models of grid cells^9,11,15^. Cross-correlograms between pairs of grid cell spike trains showed structure related to the spatial overlap between the cells’ grid fields (Figure 2). During running (Figure 2b-c, left column), grid cell pairs with highly overlapping grid fields (sorted to correspond to higher cell pair IDs in Figure 2) showed high cross-correlations between their spike trains at zero or near zero lags. Grid cell pairs with only partially overlapping grid fields tended to show moderately low correlations at short time lags, whereas grid cell pairs with anti-phase spatial tuning (corresponding to lowest cell pair IDs in Figure 2) showed the lowest correlations at short time lags. Remarkably similar correlated and anticorrelated activity patterns were observed for the same set of grid cell pairs during REM and NREM states (Figure 2a-c, middle two columns and right column). The overall widths of the short-time lag peaks and dips in the cross-correlograms were narrower during NREM than during waking and REM, suggesting that grid cell network dynamics operate on a faster (by a factor of approximately 5) time scale during NREM. This finding is consistent with temporal compression of spike sequences previously observed during NREM in networks of head direction cells^21^ and place cells^31,32^. Nevertheless, the general relationship between NREM spike-time correlations and spatial rate map correlations was qualitatively the same as that seen during waking and REM, as is particularly apparent for NREM correlations shown on a magnified time scale (Figure 2a-c, right column).

**Figure 2.**
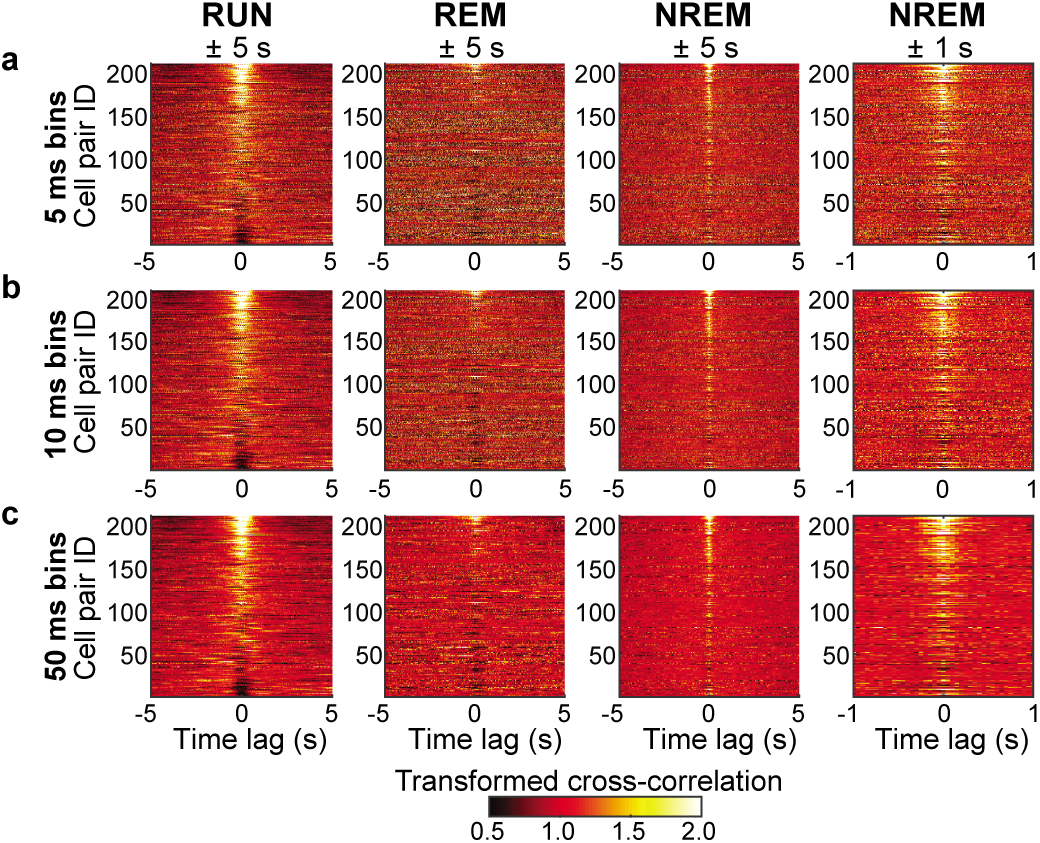
Cross-correlations between grid cell spike times were related to the degree of overlap in grid cell rate maps across active waking behaviors and sleep. (a-c) Each panel shows color-coded spike time cross-correlations (divided by their average; see Methods) for all pairs of grid cells sorted from highest rate map correlation coefficient (highest Cell Pair ID) to lowest rate map correlation coefficient (lowest Cell Pair ID) (See Methods). The leftmost three columns show results for RUN, REM, and NREM and are plotted across time lags of +/− 5 s. The rightmost column shows results for NREM plotted across time lags of +/− 1 s. Note that similar qualitative relationships between spike time cross-correlations and grid cell rate map correlations are maintained across all three behavioral states and for spike trains binned with varying degrees of temporal resolution (i.e., 5 ms bins in a, 10 ms in b, and 50 ms in c).

To quantify the qualitative trends seen in the patterns of pairwise cross-correlograms, we summed the spike time cross-correlograms within +/− 5 ms time lags to obtain a single short-time cross-correlation estimate for each grid cell pair. We then compared these estimates to their associated rate map correlations to determine the extent to which temporal correlations between grid cells’ spike trains were predicted by overlap between their rate maps across the three behavioral states (Figure 3a). Not surprisingly, grid cells’ spike train cross-correlation estimates were significantly related to their rate map correlation values during active exploration (significant regression for RUN: F (1,209) = 235.25, p < 0.0001, R^2^ = 0.53), with higher spike train cross-correlation values associated with higher rate map correlations. Remarkably though, rate map correlation values from active behaviors also significantly predicted temporal correlations between grid cells during sleep states (significant regressions for REM: F(1,209) = 52.53, p < 0.0001, R^2^ = 0.20; and NREM: F(1,209) = 93.81, p < 0.0001, R^2^ = 0.31). Therefore, despite a lack of sensory input driving the system during sleep, the relationship between cell pairs’ spike-time cross correlations and rate-map cross-correlations observed during active behaviors persisted during sleep. This finding strongly suggests that temporal correlations during both waking and sleep states result from connectivity in the grid cell network.

**Figure 3.**
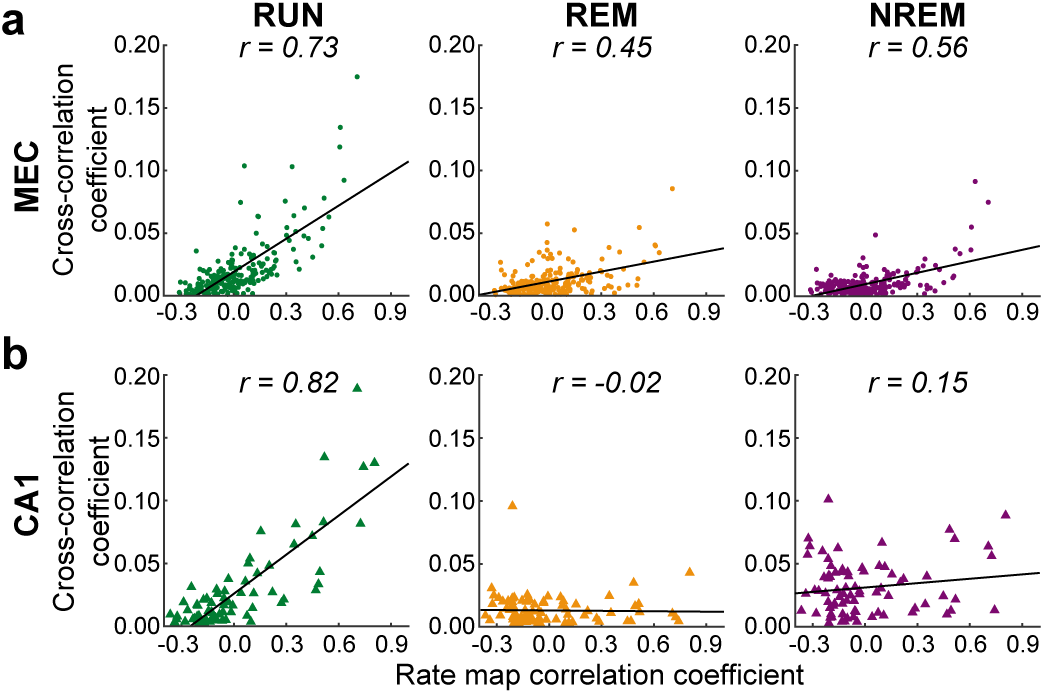
The relationship between spatial overlap and spike-time cross correlations was preserved across waking and sleep states for grid cell pairs but not CA1 place cell pairs. Spike time cross-correlation coefficients were summed across a +/− 5 ms time lag. Each dot in a scatterplot represents a grid cell pair’s rate map correlation coefficient and summed spike train cross-correlation coefficient. Each plot also shows the best-fit line in black and the associated correlation coefficient, **r**. (a) For all grid cell pairs and across all three behavioral states, positive correlations were observed between spike time cross-correlations and grid cell rate map correlation coefficients. (b) Pairs of CA1 place cells exhibited a significant positive correlation between summed spike time cross-correlation coefficients and place cell rate map correlation coefficients during active waking behaviors, but that relationship was not maintained during REM or NREM sleep.

### Preserved correlation structure in grid cells during sleep was not explained by hippocampal place cell correlation structure during sleep

With the goal of evaluating the possibility that preserved grid cell correlation relationships across awake spatial exploration and sleep were driven by place cells^33^, we next tested whether place cells’ spike time correlations during sleep were related to the overlap of their spatial fields during awake exploration. For recordings obtained during active running on a circular track (i.e., RUN), temporal correlations between 78 place cell pairs’ spike trains were significantly related to the correlation of their firing rate maps (Figure 3b (left); significant regression for RUN: F(1,76) = 160.55, p < 0.0001, R^2^ = 0.68), with temporal correlations between place cells increasing as a function of their rate map correlations. However, this relationship was not preserved in REM and NREM overnight recordings of the same place cell ensembles (Figure 3b (right panels); non-significant regressions for REM: F(1,76) = 0.04, p = 0.84, R^2^ = 0.001; and NREM: F(1,76) = 1.78, p = 0.19, R^2^ = 0.023). Accordingly, there was a significant interaction between brain region (CA1 or MEC) and behavioral state (RUN, REM, or NREM) on spike time cross-correlation values (significant multiple regression model: F(4,862) = 94.91, p < 0.001; significant interaction term: β = 0.42, p < 0.001), indicating that CA1 place cell pairs’ temporal correlations were not related to field overlap across behavioral states in the same manner as was observed for MEC grid cells. That is, the degree of place field overlap during track running only predicted the strength of place cell pairs’ temporal correlations during active waking behaviors, but not during REM or NREM sleep. Differences in relationships between MEC and CA1 field overlap and spiking co-activity across behavioral states were not explained by qualitative differences in firing rates across behavioral states. That is, CA1 firing rates between active waking behaviors and sleep followed a roughly similar pattern as MEC grid cell firing rates, with significantly lower firing rates in CA1 during REM and NREM than during active wakefulness (41 place cells; RUN: 1.61 +/− 0.25 Hz, REM: 0.85 +/− 0.14 Hz; NREM: 0.88 +/− 0.13 Hz; F(2, 120) = 5.434, p = 0.006; 1-way ANOVA; p = 0.012 for RUN vs REM, p = 0.016 for RUN vs NREM, no significant difference between REM and NREM, post hoc Tukey’s HSD tests).

### The relationship between grid cells’ relative spatial phases and spike time correlations was maintained across active and sleep states

The results shown in Figures 2 and 3 suggest that spatial map correlations are reflected, putatively through recurrent connectivity, in spike-time correlations across states. However, spike time correlations computed per cell pair are noisy, and spatial map correlations are not a perfect measure of spatial tuning relationships between pairs of grid cells. We therefore used a more detailed measure of spatial tuning relationships between cells and, to reduce noise, combined spike time cross-correlation estimates for cell pairs with similar spatial tuning offsets (see Methods). Briefly, we obtained the relative spatial phase for a given pair of cells by estimating the offset from the origin of the central peak in their spatial cross-correlogram, similar to a previous study^34^ (Figure 4a) (see Methods). This value was then used to sort cell pairs into (ΔφX, ΔφY) relative phase bins. For each cell pair, we summed the temporal cross-correlogram in a +/− 5 ms window about zero time-lag, as in Figure 3. We then averaged this value across all pairs in a bin and plotted average spike-time cross-correlations against relative spatial phase (Figure 4b, left column). This process was repeated using a broader time window of +/− 50 ms about zero in the temporal cross-correlogram to obtain a temporally coarser spike-rate cross-correlation estimate across relative spatial phases (Figure 4b, right column). We then performed the same cross-correlation computations for REM and NREM periods. To account for differing firing rates across states, we scaled each surface according to its maximum average cross-correlation value. Comparing the left and right columns of Figure 4b reveals the same general pattern at both short (spike-time) or long (spike-rate) timescales. That is, there was a strong relationship between relative spatial phase values and temporal correlations, with low relative spatial phases associated with high temporal correlations. This relationship persisted across all states regardless of whether the behavior was spatial (RUN) or not (REM and NREM).

**Figure 4.**
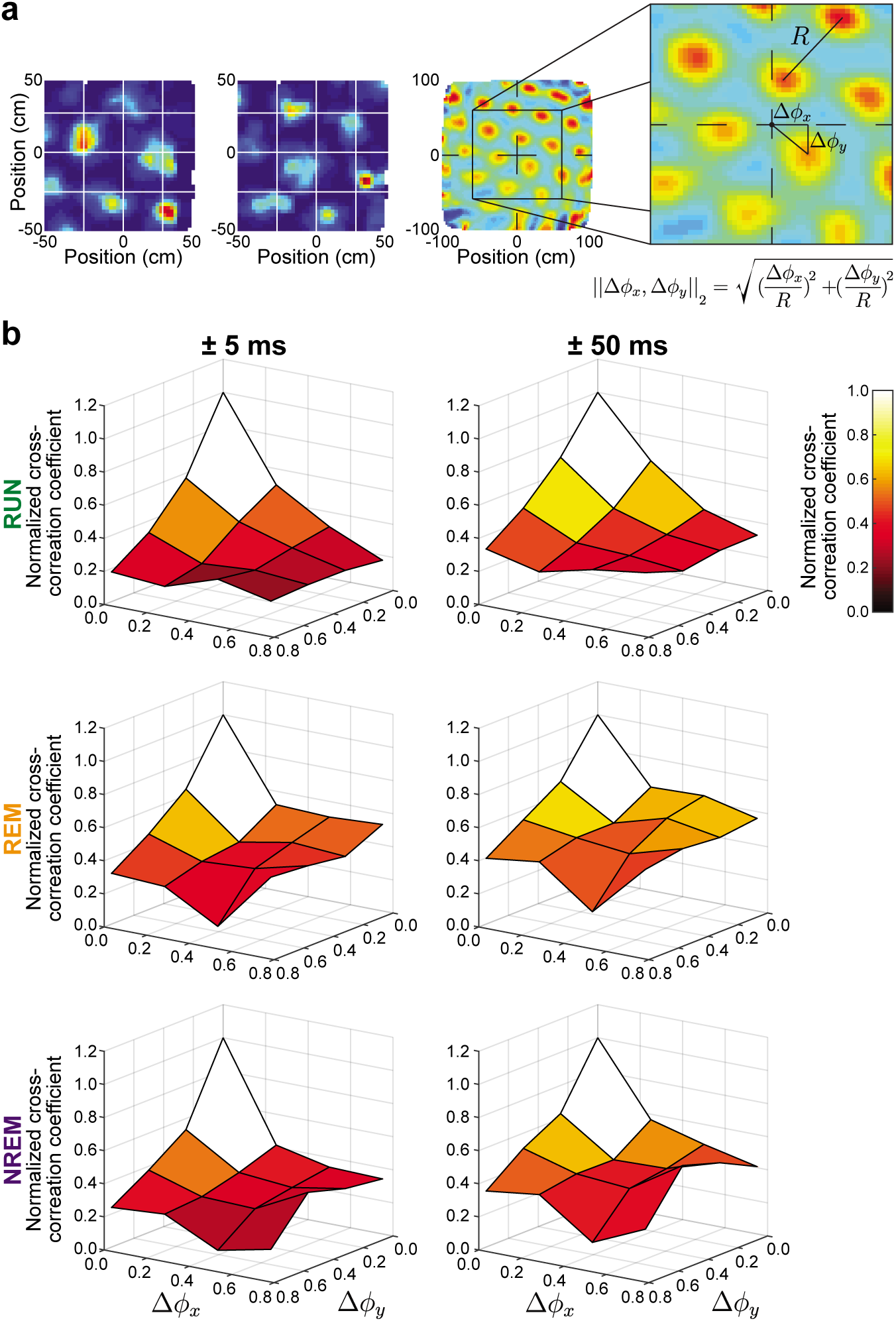
Grid cell spike time correlations during waking behaviors and sleep were predicted by spatial phase offsets between grid fields. (a) The method used to calculate the relative spatial phase between grid cells is demonstrated with this illustrative example. Color-coded rate maps for a pair of grid cells are shown in the left two panels. The third panel shows the rate map cross-correlation for the grid cell pair. The grid cell rate map cross-correlation peak nearest to the plot origin was used to estimate Δφ_X_ and Δφ_Y_ offsets (see right panel). Each offset was then normalized by the spatial period (R), which was assumed to be the same between all peaks in the rate map cross-correlation. (b) The relationship between grid cell pairs’ relative spatial phases and their spike time cross-correlations is shown. Spike time cross correlation values were summed across time lags of +/− 5 ms (left column) or +/− 50 ms (right column) and averaged within each (Δφ_X_, Δφ_Y_) bin. Data from RUN (top), REM (middle), and NREM (bottom) states show maximal spike train cross-correlation values at low relative spatial phases (i.e., high overlap of grid fields) and weaker spike train cross-correlation values at high relative spatial phases (i.e., low overlap of grid fields). The plots were scaled by their peak value to compare across behaviors with different spike rates. Normalized cross-correlation coefficient values are plotted in color scale for ease of plot interpretation.

We further compressed and reduced noise in the results by collapsing the x and y components of relative spatial phase into a one-dimensional relative spatial phase magnitude (Figure 5a for normalized cross-correlation coefficients; see Supplementary Figure 1 for non-normalized version). This transformation to one-dimensional relative spatial phase was also done to compare grid cell relative spatial phase plots to analogous metrics for CA1 place cell pairs recorded on a circular track (Figures 5b and Supplementary Figure 1). Across all three behavioral states, grid cells’ spike time correlations decreased as their relative spatial phase magnitudes increased. Temporal correlations during REM exhibited a qualitatively similar, but quantitatively weaker, functional dependence on the magnitude of relative spatial phase compared to waking and NREM states (Supplementary Figure 1 shows non-normalized plots for comparison of magnitudes). Higher noise in REM states (as is visible in Figure 2, second column) may explain this quantitative difference.

**Figure 5.**
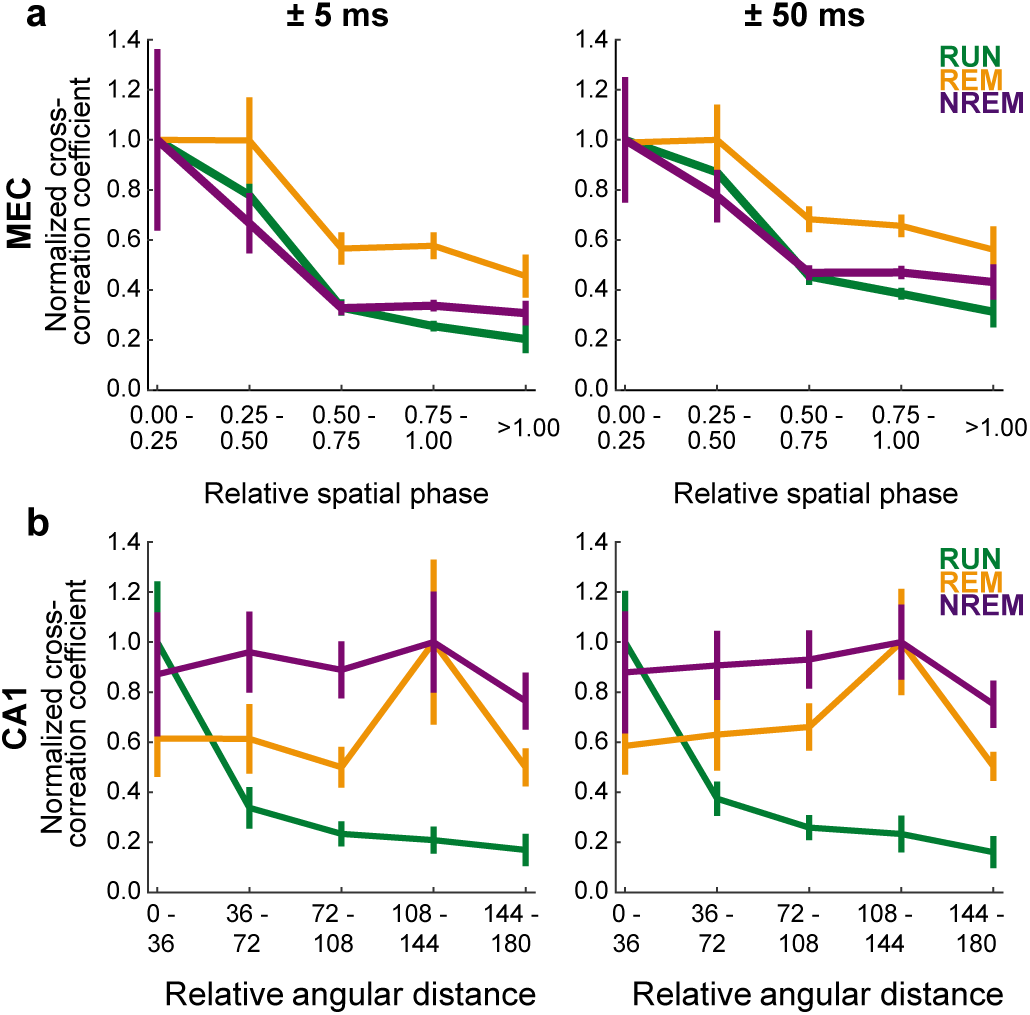
Grid cell spike time correlations decreased with relative spatial phase magnitude across behavioral states, while CA1 spike time correlations were related to distance between place fields only during active running. In this figure, spike time cross-correlation coefficients for each behavioral state were normalized by dividing by the bin with the greatest average spike time cross-correlation coefficient (see Supplementary Figure 1 for non-normalized versions). (a) In these plots, the two-dimensional relative spatial phase values depicted in Figure 4B were collapsed to a single dimension (magnitude) by calculating the Euclidean distance of Δφ_X_ and Δφ_Y_. In all three behavioral conditions, spike time cross-correlations summed over +/− 5 ms (left) and +/− 50 ms time lags (right) decreased as the magnitude of relative spatial phase increased. (b) In this plot, we calculated the relative angular distance between place field centers on the circular track and compared that distance to spike time cross-correlations summed over +/− 5 ms (left) and +/− 50 ms time lags (right). Place cells’ spike time cross-correlations decreased as distance between place fields increased during active waking behavior (i.e., RUN) but not during sleep states (i.e., REM and NREM).

In contrast to grid cells, CA1 place cell pairs’ cross-correlation coefficients during waking behaviors decreased with increasing distance between place fields, but cross-correlations during REM and NREM states did not reflect place field overlaps from the preceding spatial exploration session (Figures 5b and Supplementary Figure 1). Taken together with the results shown in Figure 3b, these results suggest that stable relationships between grid field overlap and grid cell co-activity patterns across waking and sleep states are not caused by inputs from place cells.

### Observed grid cell correlation patterns were not explained by theta coordination

The preserved relationship between grid field overlap and spike time correlations during NREM periods is noteworthy because, unlike during active wakefulness and REM sleep, theta oscillations are not present during NREM. In other words, the preserved relationship between spatial tuning and NREM coactivity patterns cannot be attributed to coordinated theta phase relationships between grid cells^35^. Moreover, spike-time correlations were higher for cell pairs with overlapping fields than pairs with largely non-overlapping fields across both theta-associated states (i.e., RUN and REM) when theta phase effects were present and when they were removed (N = 26, overlapping fields group, N = 12, non-overlapping fields; F(1,36) = 7.6, p = 0.009, repeated measures ANOVA; Supplementary Figure 2). Thus, the preservation of relationships between relative spatial phase of grid fields and spike time cross-correlations across sleep states does not appear to be explained by a shared theta modulation of cell pairs’ spikes.

### Grid cell pairs with different relative spatial phase magnitudes exhibited different distributions of spike time correlations

In addition to looking at average relationships between relative spatial phase magnitude and temporal correlations across all grid cell pairs, we categorized individual grid cell pairs according to their relative spatial phase magnitudes and plotted the associated distributions of spike time cross-correlations. We expected that the distribution of spike time correlations for cell pairs with a low relative spatial phase magnitude (i.e., high grid field overlap, example in Figure 6a) would have more mass at higher values compared to the spike time cross-correlation distribution for high relative spatial phase magnitude pairs (i.e., low grid field overlap, example in Figure 6b). Indeed, this pattern was observed when we sorted grid cell pairs into three categories according to the degree of overlap in their grid fields. Across all three behavioral conditions, the spike time correlation distributions for the low relative spatial phase magnitude group (i.e., high grid field overlap) exhibited tails containing relatively high correlation values (Figure 6c, left column). In contrast, the correlation distributions for the high relative spatial phase magnitude group (i.e., low grid field overlap; Figure 6c, right column) were more sharply peaked at lower values, across all three behavioral states. Grid cell pairs with intermediate grid field overlap showed distributions of spike time cross-correlations across all states that were intermediate between the corresponding distributions observed for high and low overlap groups (Figure 6c, middle column). Taken together with results described above, these results show that correlated grid cell activity patterns across waking and sleep reflect the degree of grid field overlap during active exploratory behaviors.

**Figure 6.**
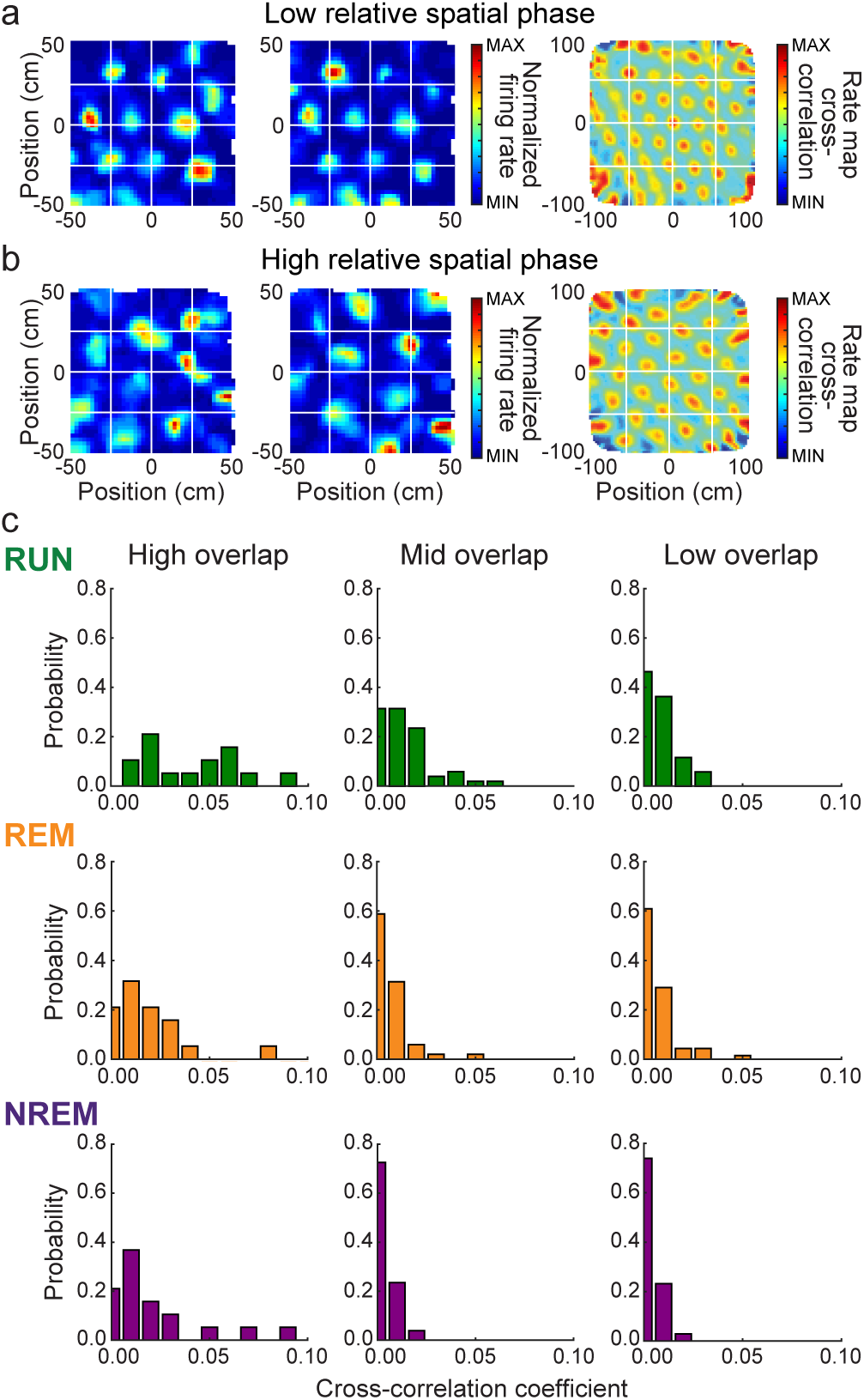
Grid cell pairs with overlapping grid fields exhibited a greater proportion of high spike time cross-correlations across active and sleep states. (a) Color-coded rate maps (left two columns) and rate map cross-correlation (right column) for an example grid cell pair with low relative spatial phase (i.e., high grid field overlap). (b) Same as A but for an example grid pair with high relative spatial phase (i.e., low grid field overlap). (c) Probability density functions for normalized spike time cross-correlation coefficients summed over +/− 5 ms time lags. Grid cell pairs were sorted into High Overlap (left column), Mid Overlap (center column), or Low Overlap (right column) categories, corresponding to low, moderate, and high relative spatial phase magnitude values, respectively. For all three behavioral conditions, grid cell pairs in the High Overlap category were more likely to exhibit relatively high spike time cross-correlations, and grid cell pairs in the Low Overlap category were more likely to exhibit low spike time cross-correlations.

## DISCUSSION

We examined whether the spatial tuning relationships of grid cells influence their spike-time correlations during non-spatial sleep states. We found that cells with strong spike-time correlations have similar spatial tuning, whereas cells with minimal spatial tuning overlap exhibit weaker or negative spike-time correlations. Further, these patterns of spatial-tuning based correlation are preserved across REM and NREM sleep states during which there is no spatial behavior. These results could not be accounted for by theta modulation, since the pattern of correlations in NREM, when theta was absent, matched that found during REM sleep and waking exploration, when theta was strong. The across-state preserved pattern of correlations as a function of spatial tuning was also not explained by hippocampal place cell inputs from CA1, since these cells did not exhibit unchanged patterns of correlation across states. Thus, the pattern of correlations is likely to originate within the entorhinal cortex rather than being inherited, through feedforward projections, from circuitry in the hippocampus. Taken together, our results suggest that recurrent connectivity is likely a major determinant of grid cell activity across awake navigation and non-spatial sleep states, consistent with continuous attractor network models of grid cells^9,11,15^. In grid cell models, the recurrent connectivity may involve both excitatory and inhibitory connections^9^ or inhibitory connections alone^9,36^. While we have seen some evidence for short-latency positive spike-time correlations between cells with high spatial overlap, with the correlation peaks preserved across states (suggesting the possibility of direct excitatory coupling between grid cells of similar phase), our evidence is not sufficient to clarify which types of connections underlie the present results, given that anatomical studies have shown both patterns of connectivity exist within the MEC superficial layers^36,37,38,39^.

It is notable that MEC grid cell correlation patterns from waking appear to be better preserved in sleep than are CA1 place cell correlations patterns. This result may seem surprising at first glance, considering that CA1 place cells replay activity patterns during sleep that resemble activity patterns from earlier waking behaviors^17,19,31,32,40^. However, our results examined correlation structure across an entire night’s sleep, while correlations between CA1 place cell spiking during active waking behaviors and subsequent NREM sleep notably decay within the first hour of sleep^17,19^. Moreover, the relationship between place cells’ field overlap and spike time correlations is stronger during awake rest than during sleep^16^. It should also be noted that different correlated spiking patterns in CA1 associated with different environments can reactivate within the same sleep episode^17^, suggesting that place cells exhibit multiple patterns of coactivation during sleep that reflect different experiences. Thus, the lack of preserved correlated activity between CA1 place cells across sleep states is consistent with higher-dimensional dynamics in the hippocampus during waking, which allow for globally remapped representations of different environments^41^. It is possible that hippocampal sleep states explore this higher-dimensional space, activating multiple representations that reflect waking trajectories from a variety of different environments. By contrast, the states in one grid module maintain the same low-dimensional dynamics across non-active and sleep states that they exhibit during awake exploratory behaviors in novel and familiar environments of different shapes and configurations^23,34,42^. This rigid confinement of the dynamics of grid cells in a module, across behaviors and states, implies and predicts that the grid cells will likely exhibit the same preserved cell-cell relationships during other modes of behavior, such as during navigation through non-spatial and other conceptual spaces^29,30,43^. This prediction is readily testable, using simultaneously recorded co-modular grid cells. Indeed, all models of grid cells that are based on integrating a velocity signal to generate an updated estimate of grid phase (these include continuous attractor models and oscillatory interference models^10,12,13,28^) are applicable to navigation through non-physical continuous spaces, so long as the velocity input to the grid cells reflects the time-derivative of the corresponding continuous variable. However, the predictions of preserved low-dimensional structure through conserved cell-cell relationships during non-spatial navigation follow specifically from low-dimensional attractor models^9,28,34,44^.

Within the class of continuous attractor models are networks with different types of recurrent connectivity and topology. One possibility is that the grid cell network supports the activation of one local group of interacting neurons (an activity bump) in the neural sheet and that connectivity is global (single-bump networks^45^). Another is that the connectivity is local (according to some proper rearrangement of neurons) and that activity consists of multiple activity bumps on the cortical sheet^9,28^. In the former network, co-active neurons are recurrently coupled and should always show high correlations. In the latter, co-active neurons in different bumps may share no direct coupling, and thus the distribution of correlations given the same tuning phase should be bimodal. The slight hint of bimodality in the correlations of cells with high spatial tuning overlap (far left column in Figure 6c, particularly during RUN) may be suggestive of a multi-bump network. However, additional experimentation with circuit perturbation techniques will be required to determine which connectivity pattern most accurately describes the system^46^.

The finding that grid cell co-activity during sleep resembles co-activity during waking behaviors raises the possibility that superficial-layer MEC grid cells play a role in offline memory processing. This hypothesis is consistent with reports involving recordings from other cortical areas. For example, neuronal activity in primary visual cortex^47^, auditory cortex^48^, and medial prefrontal cortex^49^, as well as grid cell activity in deep layers of MEC^50^, has been reported to correlate with hippocampal place cell activity during replay events, which are hypothesized to play a role in memory consolidation. However, the present results only involve grid cells recorded from MEC superficial layers. An initial study of grid cell replay in MEC superficial layers reported that grid cells rarely show coordinated reactivation with hippocampal place cells^26^, a result that seems inconsistent with the hypothesis that superficial layer grid cells drive consolidation of hippocampal memory representations during sleep.

The present results emphasize a need for caution when searching for evidence of replay in superficial-layer grid cells during sleep. That is, the extent to which replay occurs in the superficial layer grid cell network may be overestimated due to the structured nature of instantaneous activity patterns that reflect stable, long term network connections rather than plasticity induced by earlier experiences. Investigations of replay in superficial layer grid cells would therefore be strengthened by the inclusion of additional behavioral controls, such as comparisons of co-activity patterns during sleep to co-activity patterns during subsequent exploration of a novel environment. Grid cell simulations would also be helpful to determine the extent to which it is possible to distinguish between co-activity due to long-term network connections versus more recent experience-dependent reactivation. Such simulations would likely provide useful information about the number of simultaneously recorded superficial layer grid cells necessary to reliably detect experience-dependent reactivation. Future experiments, designed to determine whether sleep-associated reactivation of correlated firing patterns in ensembles of superficial layer MEC grid cells is experience-dependent, are key to achieving a deeper understanding of neural and network mechanisms of memory consolidation in the entorhinal-hippocampal network during sleep.

## ONLINE METHODS

### Subjects

MEC and CA1 data were collected from six male Long-Evans rats weighing ∼450 g to ∼650 g (mean +/− std = 549 +/− 87 g) and four male Long-Evans rats weighing ∼400g to ∼530g (482 +/− 55 g), respectively. After surgery, animals were individually housed with several items for enrichment (e.g., cardboard tubes, wooden cubes, plastic toys). Rats were housed in custom-built acrylic cages (40 cm x 40 cm x 40 cm) on a reverse light cycle (Light: 8pm to 8am). Active waking behavior recordings were performed during the dark phase of the cycle.

Rats recovered from surgery for at least one week, with free access to food, before behavioral training resumed. During the training and data collection period, rats were food-deprived to no less than 90% of their free-feeding weight. During periods of food deprivation, food was provided ad libitum for one day per week. All experiments were conducted according to the guidelines of the United States National Institutes of Health Guide for the Care and Use of Laboratory Animals under a protocol approved by the University of Texas at Austin Institutional Animal Care and Use Committee.

### Recording drive

For animals receiving MEC recording implants, chronic “hyperdrives”^51^ (5 animals) or a Harlan Drive (Neuralynx, Bozeman, Montana, 1 animal) with 12 tetrodes and two reference electrodes or 16 tetrodes, respectively, were implanted over MEC in the right hemisphere each animal. CA1 animals were implanted with hyperdrives above the hippocampus in the right hemisphere. Tetrodes were constructed from 17 μm polyimide-coated platinum-iridium (90 / 10%) wire (California Fine Wire, Grover Beach, California). Electrode tips of tetrodes designated for single unit recording were plated with platinum to reduce single channel impedances to ∼150 to 300 kOhm at 1 kHz.

### Surgery

Anesthesia was induced by placing an animal in an induction box filled with isoflurane (5%) mixed with oxygen (1.5 liters per minute). The animal was then moved to a stereotaxic frame, and isoflurane anesthesia (∼1.5-3%), mixed with oxygen as above, was administered throughout surgery. Buprenorphine (0.04 mg / kg) was administered subcutaneously prior to the initial incision. Subjects also received 0.04 ml of atropine (0.54 mg/ml) injected subcutaneously prior to surgery and an additional 0.02 ml of the atropine solution after four hours of anesthesia to prevent fluid accumulation in the lungs. Sterile saline (1.00 ml, 0.90%) was administered subcutaneously prior to surgery and each hour thereafter for hydration. Subjects were checked for breathing rate and responsiveness every 15 minutes during surgery to monitor anesthesia levels.

Seven to eight bone screws were placed in the lateral, anterior, and posterior edges of rats’ skulls, serving as anchors for the recording drives. Two additional screws in the anterior portion of the skull, ipsilateral to the recording drive, were used as the electrical ground reference for recording. For MEC implants, drives were positioned 4.5 mm lateral from the midline and approximately 0.2 mm anterior to the transverse sinus. The most posterior tetrode was used to position the drive according to these coordinates. For CA1 implants, drives were centered at 3.0 mm lateral to the midline and 3.8 mm posterior to bregma. Silicon adhesive (Kwik-Sil; WPI, Sarasota, Florida) was used to fill in the exposed portion of the craniotomy, and dental cement was applied to anchor the drive to the skull and to the bone screws. At the end of surgery, subjects were given a subcutaneous injection of Rimadyl (5 mg / kg), mixed with ∼1 ml of sterile saline (0.9%), for pain management.

### Data collection

Data were recorded using a Digital Lynx system and Cheetah 5.0 recording software (Neuralynx, Bozeman, Montana). Local field potentials (LFPs) from one channel within each tetrode were continuously recorded at 2 Khz, using the built in high-pass (DC-offset filter at 0.10 Hz with 0 taps) and low pass (FIR filter at 500 Hz using 64 taps) filters. Input amplitude ranges were adjusted before each recording session to maximize resolution without signal saturation. Input ranges generally fell within +/− 1600 to +/− 2600 μV. Spikes were detected and recorded in the following manner. All four channels within each tetrode were filtered from 600-6000 Hz using a high pass (FIR filter at 600 Hz using 64 taps) and low pass (FIR filter at 6000 Hz using 32 taps) filter. Spikes were detected when the filtered continuous signal crossed a threshold set daily by the experimenter, which ranged from 40-75 μV. Detected events were acquired with a 32 KHz sampling rate. Signals were recorded differentially against a dedicated reference channel (see “Tetrode Positioning” section below).

Video was recorded through the Neuralynx system with a resolution of 720 x 480 pixels and a frame-rate of 29.97 frames per second. Animal position and head direction were tracked via an array of red and green LEDs on an HS-54 headstage (Neuralynx, Bozeman, Montana) attached to a hyperdrive, or red and green LEDs connected to EIB-36-24TT headstages (Neuralynx) attached to a Harlan Drive.

### Tetrode positioning

All tetrodes were initially lowered ∼900 μm on the day of surgery. Thereafter, tetrodes were lowered gradually over the course of several weeks to the superficial layers of MEC or dorsal CA1 guided by the estimated depth and known electrophysiological hallmarks of MEC (e.g., prominent theta oscillations)^35,52^ and CA1 (e.g., sharp-wave ripples)^53^. In the CA1 implanted rats and all but one of the MEC implanted rats, the most anterior tetrode was designated as the reference for differential recording and was targeted toward the angular bundle (MEC animals) or a quiet region of overlying cortex (CA1 animals). Due to prominent noise on the most anterior channels in one MEC rat, the most posterior tetrode was chosen as a reference instead. Tetrode positions were verified histologically after experiments were completed (see Histology section below). Reference tetrodes were continuously recorded against ground to ensure that they remained in electrically quiet locations across all days of recording. Experimental sessions and data recording began when the presence of at least four simultaneously recorded grid cells was detected (see Grid Cell Detection section below) or, in the case of CA1 recordings, when most of the tetrodes were estimated to be in the CA1 cell body layer.

### Behavioral training of MEC rats

The MEC implanted rats were pre-trained, prior to surgery, to run in multiple environments, including a 1 m x 1 m open field, for randomly scattered small pieces of sweet cereal or cookie rewards. Once rats explored the open field relatively uniformly with few pauses, chronic recording drives were surgically implanted (see Surgery section above). After recovery from surgery (i.e., approximately one week after implantation), rats were familiarized over the course of three days with an open field arena (1 m x 1 m with 0.5 m wall height) in a recording room, in which they again foraged for small pieces of cereal or cookie. Open field familiarization days consisted of alternating periods of three open field sessions (20 min each), with rest sessions (10 min each) preceding and following each active session. During each rest session, rats were placed in a towel-lined, elevated flower pot. Following these familiarization days, open field foraging behavior was maintained with daily training consisting of two to three 20-minute open field foraging sessions, each preceded and followed by 10-minute rest sessions. When prominent saw-toothed theta oscillations (6-10 Hz) were detected on the recording tetrodes, suggesting that tetrodes had reached MEC, open field foraging continued to be maintained as described above, except with 30-minute, rather than 10-minute, intervening rest sessions.

On days when sufficient numbers of simultaneously recorded putative grid cells were observed, data collection began. On these days, rats also ran on a linear track after their open field recordings were completed and approximately 1-2 hours before their overnight sleep recordings began. These linear track data were not included in the present study but have been used in other studies^54,55^. At the beginning of their light cycle (∼8 pm), rats were placed in a relatively small open field arena (60 cm x 60 cm) for overnight sleep recordings. In this small arena, rats were provided with cloth bedding and toys for enrichment, as well as access to their food and water. At approximately 8 am the following morning, rats were returned to their home cages. Following ∼2-3 hours of rest in their home cages, rats were returned to the larger (i.e., 1 m x 1 m) open-field arena for an abbreviated recording session (two 20-minute bouts of open field foraging, with three intervening 10-minute rest bouts). This second open field recording was performed to confirm the stability of overnight grid cell recordings (see Grid Cell Detection section below).

### Behavioral training of CA1 rats

Rats implanted with drives in CA1 were initially part of a different study, and thus different behavioral training protocols were used. Prior to surgery, CA1 implanted rats were trained to run on a circular track with a diameter of 100 cm and a track width of 10 cm. Small pieces of sweet cereal were placed on opposite ends of the circular track as food reward. Once the rats reliably ran at least ten laps on the track in a 10-minute session, hyperdrive recording devices were implanted. Following one week of recovery, rats were re-trained to run on the circular track. Each training day consisted of three 10-minute running sessions interleaved with four 10-minute rest sessions. All CA1 implanted rats received a minimum of three days of training on the circular track before data collection began.

On each recording day, CA1 implanted rats ran on the circular track for four 10-minute running sessions with intervening 10-minute rest sessions. Upon completion of the circular track task (∼8pm), rats were placed in a 60 cm x 60 cm open field enclosure, with water and food provided along with cloth bedding and enrichment items, for overnight sleep recordings.

### Histology

Histological sections were prepared in the following manner to allow for verification of tetrode positions after completion of recordings. Rats were given a lethal i.p. injection of Euthanasia III solution. This was followed by transcardial perfusion, first with physiological saline to clear blood and then with 4% formaldehyde solution to fix brain tissue. Brains from MEC-implanted rats were cut sagittally at 30 μm, and each section through the relevant portion of MEC was collected. Brains from CA1 implanted rats were cut coronally at 30 μm, and each section through the relevant portion of CA1 was collected. Sections were mounted on slides and stained with cresyl violet. Tetrode tips and tracks were localized by comparing across successive sections.

### Spike sorting

MCLUST cluster-cutting software (version 3.5; A.D. Redish, University of Minnesota, Minneapolis) run in MATLAB 2014a (The MathWorks, Inc., Natick, Massachusetts) was used to sort spikes into putative single units. Spike waveforms were sorted based on peak height, waveform energy, and peak-valley difference. The valley depth of spike waveform was used as an additional feature to sort MEC units. A putative single-unit was accepted for further analysis if the associated cluster had a minimum 1 ms refractory period and shared less than 1% of its total number of spikes with any other accepted cluster.

### Identification of active waking and quiescent sleep states

For the initial identification of sleep epochs, videos of rats’ overnight behavior were manually scored by two independent researchers trained to discriminate between sleep, stationary alertness, and consummatory behaviors. Only epochs that both researchers classified as sleep were accepted for further analysis. REM and NREM epochs were then classified during identified sleep epochs based on oscillatory activity in the LFP using the following criteria. A recording period was classified as NREM if the moving window average (5.0 s window, 0.5 s step) of the theta (6-10 Hz) power to delta (2-5 Hz) power ratio remained below 1.0 for at least 15 seconds (modified from a previous study^56^). Instantaneous theta and delta power were calculated via Morlet wavelet transform^57,58^ calculated with a width parameter (**σ**) of 5 and a frequency resolution of 1 Hz. Recording periods were classified as REM if the moving window average of the theta-delta power ratio remained above 1.5 for at least 60 seconds (modified from previous studies^40,56^).

Recordings of active waking behaviors on the circular track (CA1 data) and in the open field (MEC data) were used to define periods of active ambulation (i.e., RUN). RUN epochs were defined as periods in which the theta-delta power ratio was above 2.0 and a rat’s running speed, smoothed with a Gaussian window (radius of 133 ms or 4 video frames, standard deviation of 1), was greater than 5 cm / sec (modified from a previous study^56^). Thresholds were confirmed by manual examination of the power spectra and randomly chosen segments of LFP to confirm the presence or absence of theta oscillations.

### Grid cell detection

Recordings of single unit activity in the open field environment were used to detect grid cells. First, rate maps were calculated using 3 cm spatial bins and smoothed with a Gaussian window^1^. A gridness score^3^ was then calculated for each putative grid cell and compared to a bootstrapped distribution of gridness scores. The bootstrapped distribution consisted of 2000 shuffles in which spike trains were shifted in time by a random amount while the inter-spike intervals remained fixed. Rate maps and gridness scores were recalculated for each shuffle to create a bootstrap distribution for each putative grid cell. Units were classified as grid cells when the original gridness score met or surpassed the 95th percentile of the corresponding bootstrapped distribution. Additionally, a grid cell was only included if the same unit was identified and remained stable in an open field recording session the next day. Grid cell stability was assessed by visual inspection of unit cluster position, inter-spike intervals, and grid field similarity.

### Grid cell firing rate estimation

Grid cell firing rates during each RUN, REM, and NREM period were calculated by dividing the total number of spikes by the duration of the detected behavioral period, averaged across periods for each behavioral state (Figure 1c, histograms), and then averaged across units (Figure 1b).

### Place cell firing rate map calculation

Given that CA1 place cells were recorded while rats performed laps around an elevated circular track, rather than ambulating throughout an open field environment as with grid cell recordings, the procedure for rate map calculation was different for place cells than for grid cells. For CA1 recordings on the circular track, rats’ positions were first converted from two-dimensional Cartesian coordinates to a one-dimensional measure of angular position along the track. The sequence of angular coordinates corresponding to a rats’ position at each video frame time were then sorted into 5-degree bins spanning from 0 to 360. Rate maps were calculated by dividing the total number of spikes occurring while the rat was actively ambulating within each radial bin by the total amount of time that the rat spent actively moving within that bin. They were then smoothed with a Gaussian kernel 25 degrees wide.

### Statistics

Unless otherwise noted, all analyses were performed in MATLAB 2017a (The MathWorks, Inc., Natick, Massachusetts) using custom-written software. However, standard built-in MATLAB functions were used (e.g. “xcorr,” “corrcoeff,” etc.) whenever possible.

The SPSS Statistics Subscription (build 1.0.0.781, IBM) software package was used for the statistical analyses in this paper. This includes the statistics describing firing rate differences across behaviors (as shown in Figure 1 for MEC), the statistics comparing the relationships between rate map correlation coefficients and the spike time correlation values (calculated from the sum of the correlations within the +/− 5 ms lag window) for data in Figure 3, as well as the statistics used to analyze the effects of removing theta phase influences (Supplementary Figure 2).

### Spike time correlations

Spike time cross-correlations were calculated in MATLAB using the “xcorr” function (MATLAB 2017a Signal Processing Toolbox, The MathWorks, Inc., Natick, Massachusetts). Unless otherwise stated, cross-correlations were normalized such that auto-correlations at zero lag were identically 1.0 (i.e., “coeff” normalization option in MATLAB), and correlograms were averaged across bouts of the same behavior. The exception to this is Figure 2, in which each of the un-normalized cross-correlograms, which were summed across bouts of the same behavior, were scaled by their average value (Gardner et al., SfN Abstracts 2016). This analysis was done to compare the present results with those of Gardner and colleagues, who introduced this analysis previously. Cross-correlations were calculated using spike counts in 2 ms (Figures 3-6, Supplementary Figure 1), or 5 ms, 10 ms, and 50 ms bins (Figure 2a, b, and c, respectively) to ensure that the observed results were not an artifact of the chosen bin size. All cross-correlations were calculated over +/− 5000 ms lags. MEC and CA1 cell pairs that did not have a peak of at least 5 spikes in the (summed un-normalized) cross-correlation for each behavioral state were not considered. MEC cell pairs recorded simultaneously on the same tetrode were excluded if spike time correlations at zero lag were greater than the average of the correlation vector to prevent double counting of the same spikes. MEC cell pairs in which the spatial period of one cell was not within 30% of the other were also excluded to restrict comparisons to cells within the same putative module^34^.

### Rate map correlation coefficients

Correlation coefficients for pairs of grid cell rate maps, which were used to sort grid cell pairs in Figure 2 and are shown across the x-axes in Figure 3a, were calculated by the off-diagonal value returned by the “corrcoef” function in MATLAB (MATLAB 2017a, The MathWorks, Inc., Natick, Massachusetts). The values presented are the medians across the three open-field sessions. For CA1 place cells (Figure 3b), the rate map correlation coefficient was calculated using the ‘corrcoef’ function, using rate maps that were constructed from angular positions on the circular track.

### Grid cell rate map cross-correlations

Rate map autocorrelations (for grid cell detection) and cross-correlations (Figures 4, 5a, 6; Supplementary Figure 1) were performed following procedures outlined previously^1^. Briefly, the two-dimensional cross (or auto) correlation was calculated from the smoothed rate maps via:

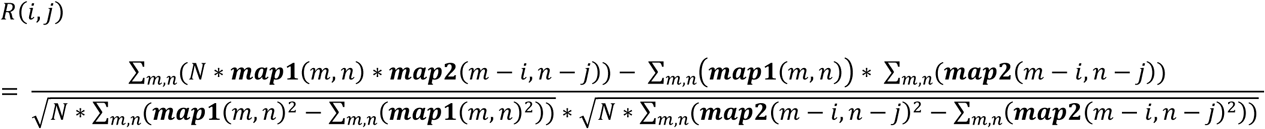

Where **map1** and **map2** are the smoothed firing rate maps and N is the total number of spatial bins.

### Relative spatial phase between pairs of grid cells

Relative spatial phase was calculated using the rate map cross-correlation between the rate maps of two grid cells (Figures 4, 5a, 6; Supplementary Figure 1; see Figure 4a for a schematic explanation of the method) (similar to a previous study^34^). The peak of the cross-correlation map closest to the plot origin was identified, and spatial phase was calculated using the x and y distances from the origin to that peak. The x and y distances were then each normalized by the spatial period, calculated as the distance between the two peaks closest to the origin. Because the distance of least overlap is at one half the spatial period, phases greater than 0.5 were subtracted from 1.0 (e.g., 0.62 becomes 1.00-0.62 = 0.38). The phase was then divided by 0.5, giving a final relative spatial phase value that ranged from [0, 1] where [0, 0] indicated maximal overlap between grid fields and [1, 1] indicated minimal overlap between grid fields. The medians of the relative phase values calculated across the three open field sessions were used as the final relative spatial phase values shown in Figure 4. A one-dimensional measure of the relative spatial phase magnitude was also created by taking the Euclidian distance of the x and y relative phases, which spanned the range [0, 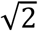] (Figures 5a, 6; Supplementary Figure 1, Supplementary Figure 2).

Cell pairs in Figure 6 were categorized as “High Overlap” (i.e., low relative spatial phase), “Low Overlap” (i.e., high relative spatial phase), or “Mid Overlap” (i.e., intermediate relative spatial phase) by comparing each cell pair’s relative spatial phase magnitude to the maximum observed relative spatial phase magnitude (which could range in value from 0 to 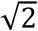). Cell pairs with a relative spatial phase magnitude that fell below 30% of the maximum relative spatial phase magnitude were labeled as High Overlap. Cell pairs with a relative spatial phase magnitude that fell above 70% of the maximum relative spatial phase magnitude were classified as Low Overlap. Cell pairs were assigned to the Mid Overlap group if their relative spatial phase magnitude fell between 40% and 60% of the maximum relative spatial phase magnitude. Cell pairs not assigned to one of these three groups were excluded from results shown in Figure 6. Supplementary Figure 2 used the same criteria for the High Overlap and Low Overlap groups, but did not include a Mid Overlap group.

### Relative distance between pairs of place cells

The relative angular distance between the firing locations of two CA1 cells on the circular track was defined as the length of the minor arc between their fields. Only place cells with peak firing rates exceeding 1 Hz were included in this analysis. Each cell pair was sorted into one of five bins based on the relative angular distance (each bin spanned a range of 36 degrees), and the average spike time correlation across all pairs within a group was calculated (Supplementary Figure 1). Similar to Figure 5a, spike time correlations for each behavioral state were then normalized by the maximum average spike time cross-correlation coefficient for that behavioral state (Figure 5b).

### Removal of theta contribution to spike time cross – correlations

One possible explanation for the relatively high spike time cross-correlations observed for some grid cell pairs, aside from network connectivity, is that spikes from those grid cells tended to occur at the same theta phase. To address this possibility, the observed spike time cross-correlations in each behavioral period (RUN or REM) were compared to distributions containing 200 spike-time cross-correlations computed from shuffled spike train pairs, based on a method reported previously^59^. In these shuffled spike trains, the times of each spike for each grid cell were shuffled independently. Each spike was moved randomly to a new time that satisfied two conditions: the new time fell within a +/− 500ms window around its original spike time and was associated with a theta phase within 5 degrees of the original spike time’s associated theta phase. Thus, theta-induced correlations and other correlations on a time-scale slower than 500 ms in the shuffled data were preserved, while short-latency correlations were eliminated. Examples of individual cross-correlations for shuffled spike trains are shown in Supplementary Figure 2 together with the corresponding original cross-correlation. The average of the cross-correlations for all shuffled spike trains was presumed to represent the portion of the cross-correlation explained by theta phase co-modulation alone (Supplementary Figure 2). To visualize the temporal correlations of spikes that were not attributable to theta phase modulation of spike times, the magnitude of the observed cross-correlation that fell within the 98% confidence interval of shuffled distribution was reduced by the amount attributable to theta or other slow correlations (Supplementary Figure 2). To do so, first, any lag bins of the original correlogram with values that fell within the confidence interval were set equal to zero. Next, in any time lag bins in which the correlogram was greater than the upper bound of the confidence interval, the correlogram was reduced by the value of the upper confidence interval bound in that bin (e.g. if the correlogram in a bin was equal to 0.1 and the upper confidence interval bound was 0.06, the new correlogram value would be 0.04). Values of the correlogram which fell below the lower bound of the confidence interval were reduced by the value of the lower bound (e.g. if the correlogram bin was equal to 0.02 and the lower confidence interval bound was 0.03, the new correlogram value would be-0.01).

For each cell pair, the average theta-removed cross-correlation across all sessions of a given behavior was calculated by taking the theta-removed cross correlation for each individual behavioral session and then averaging across sessions. The area of the cross-correlogram between +/− 5 ms lag was calculated (“summed correlation”, as in Figures 3, 4B left, 5 and 6C and Supplementary Figure 1) and used as a measure of short-latency correlation in Supplementary Figure 2. The population average of the summed correlation was used in Supplementary Figure 2 to look at differences between low and high relative spatial phase groups, calculated as in Figure 6. The relationship between relative spatial phase and the summed correlation or the summed, theta-subtracted correlation (Supplementary Figure 2) was tested using repeated measure ANOVA.

## Data availability

The data used in this experiment is available upon request from the authors.

## Code availability

The custom MATLAB scripts used in this paper are available upon request from the authors.

## ACKNOWLEDGEMENTS

We thank K.N. Bobbitt and K. Kallina for recording drive construction and outstanding technical support. We also thank A. Akinsooto, K.N. Bobbitt, S. Brizzolara-Dove, J. Campos, A. Davis, D. Jones, G. Kwong, C.G. Orozco, F. Rahman, E. Usheva, and D. Wehle for help with sleep video scoring, and B.J. Gereke for helpful discussions. This work was supported by: the Whitehall Foundation (to L.L.C.), NSF CAREER Award 1453756 (to L.L.C.), ONR YIP Award N00014-14-1-0322 (to L.L.C.), and the National Institute on Drug Abuse (primary) of the National Institutes of Health under award number T32DA018926 (to E.H. and S.T.). I.R.F. was supported in part by a Faculty Scholar award from the Howard Hughes Medical Institute, a grant from the Simons Foundation through the SCGB, and an HFSP grant.

## AUTHOR CONTRIBUTIONS

S.G.T., I.F., and L.L.C. designed experiments and analyses. S.G.T. and E.H. collected data. S.G.T., J.B.T., and E.H. wrote analysis programs. S.G.T., E.H., and J.B.T. analyzed data. I.F. and L.L.C. supervised the research. S.G.T., J.B.T., I.F., and L.L.C. wrote the paper, with comments from E.H. All authors discussed results.

## FIGURE LEGENDS

**Supplementary Figure 1.**
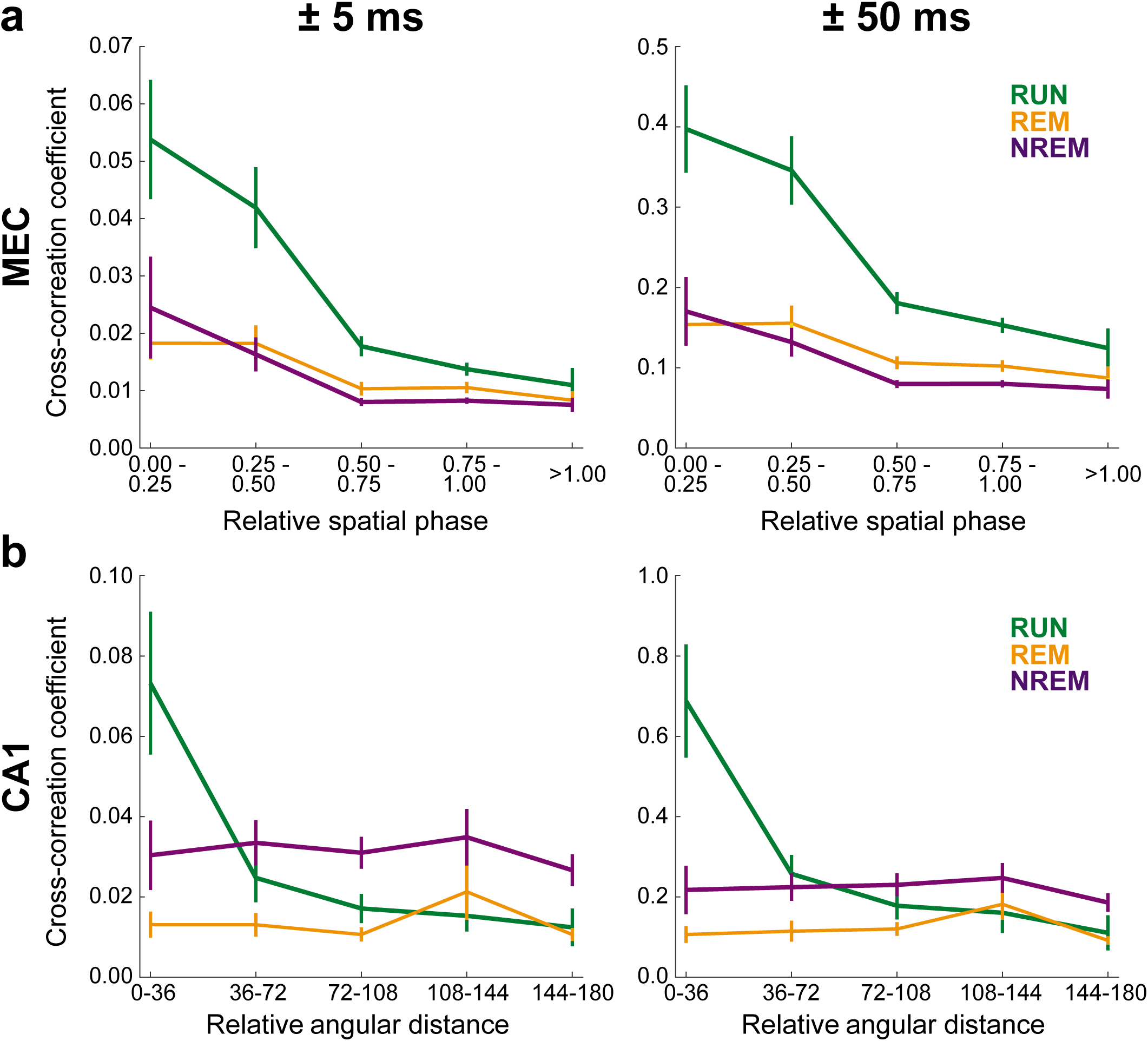
Related to Figure 5; Across all behavioral states, grid cell pairs’ cross-correlation coefficients decreased with increasing relative spatial phase. Place cell pairs’ cross-correlation coefficients decreased with increasing distance between place fields during RUN, but a similar relationship was not observed during NREM or REM. These plots are the same as those in Figure 5 except that spike time cross-correlation coefficients were not normalized according to maximum values within each behavioral state.

**Supplementary Figure 2.**
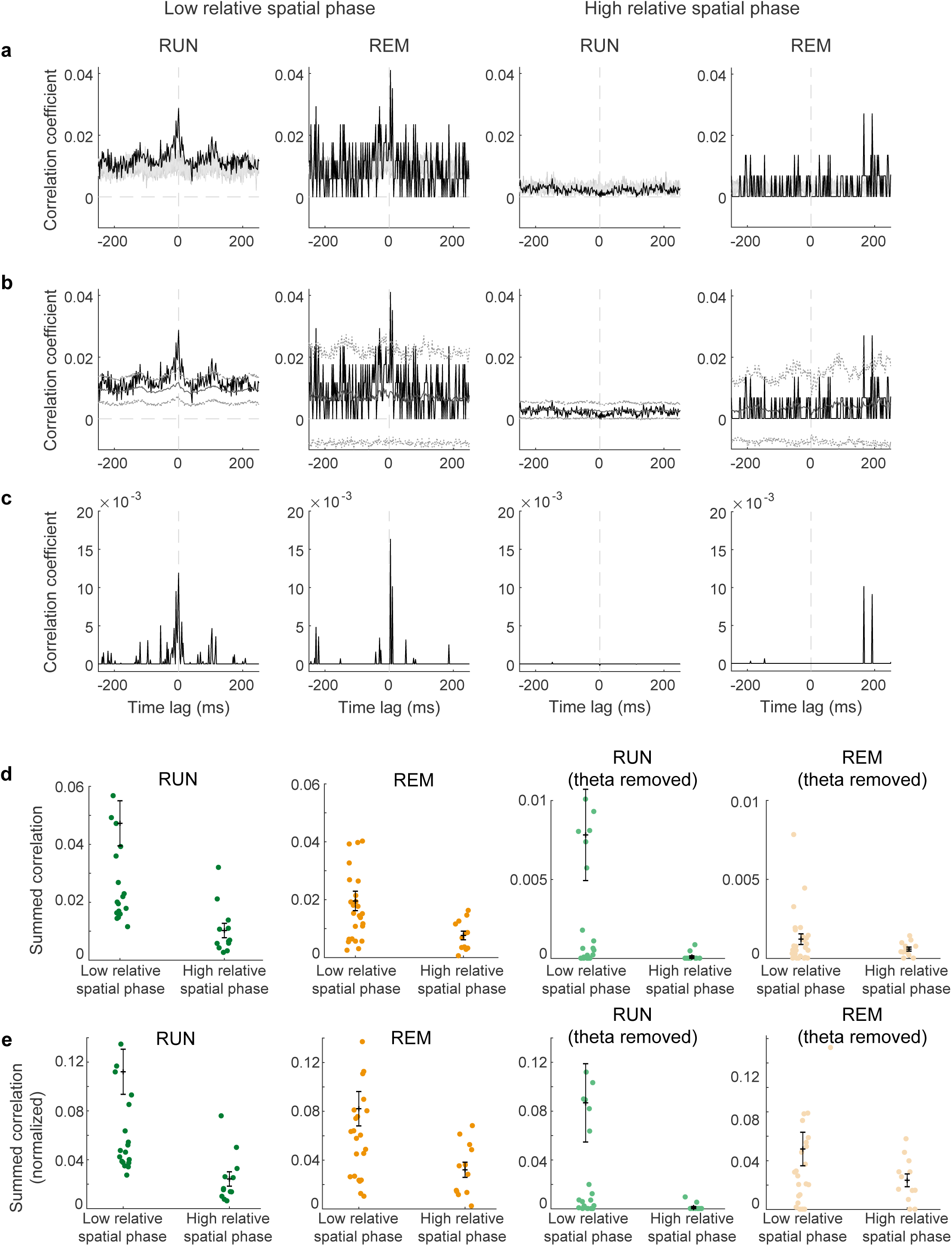
Related to Figure 5; Short-time spike time correlation patterns of grid cell pairs with overlapping grid fields maintained across behavioral states when the effect of theta phase modulation of spiking and other slow influences was removed. To determine the extent to which spike time cross-correlation results were explained by shared theta phase preferences of spike times and other slow modulations in spike rate, spikes were temporally shuffled two hundred times within 500 ms moving windows, while keeping the theta phase of each spike time fixed. Spike time cross-correlations were then re-calculated using shuffled trains (See Methods). This analysis was only done for RUN and REM spike trains, given that NREM is characterized by an absence of theta. (a) Shown are original spike time cross-correlations (black) overlaid on 10 (out of 200) randomly selected examples of shuffled cross-correlations (gray) for an example grid cell pair. Cross-correlograms for an example grid cell pair with highly overlapping fields (i.e., low relative spatial phase magnitude) are shown for RUN (first column) and REM (second column). Also shown are RUN (third column) and REM (fourth column) cross-correlograms for an example grid cell pair with largely non-overlapping grid fields (i.e., high relative spatial phase magnitude). (b) The average theta-determined component of the correlation (gray, solid) with 98% confidence intervals (gray, dotted) and associated original spike time cross-correlogram (black). Columns are as in A. (c) The correlogram remaining after removal of the theta-modulated components. Note that substantial peaks around zero lag were still present in the low relative spatial phase plots (left two columns) but not in the high relative spatial phase plots (right two columns). (d) Cell pairs were split into low and high relative spatial phase groups as in Figure 6, and their correlation was summed over the +/− 5 ms lag window. The average correlation sum was calculated across each subpopulation for both RUN (first column) and REM (second column). The same calculation was done on the theta-removed correlations in the two columns to the right. (e) The same as in D, but with values normalized by the Euclidean norm of the population of summed correlations before sorting into near or far groups. This was done to aid visual comparison between RUN and REM conditions, and between original and theta-removed groups.

## References

1. Hafting, T., Fyhn, M., Molden, S., Moser, M.-B. & Moser, E. I., Microstructure of a spatial map in the entorhinal cortex. Nature 436, 801–806 (2005).

2. O’Keefe, J. & Dostrovsky, J., The hippocampus as a spatial map. Preliminary evidence from unit activity in the freely-moving rat. Brain Research 34, 171–175(1971).

3. Sargolini, F. et al., Conjunctive representation of position, direction, and velocity in entorhinal cortex. Science 312, 758–62 (2006).

4. Taube, J. S., Muller, R. U. & Ranck, J. B., Head-direction cells recorded from the postsubiculum in freely moving rats. I. Description and quantitative analysis. The Journal of Neuroscience 10, 420-35 (1990).

5. Solstad, T., Boccara, C. N., Kropff, E., Moser, M.-B. & Moser, E. I., Representation of geometric borders in the entorhinal cortex. Science 322, 1865–8 (2008).

6. Kropff, E., Carmichael, J. E., Moser, M.-B. & Moser, E. I., Speed cells in the medial entorhinal cortex. Nature 523, 419–424 (2015).

7. Hardcastle, K., Maheswaranathan, N., Ganguli, S. & Giocomo, L. M., A Multiplexed, Heterogeneous, and Adaptive Code for Navigation in Medial Entorhinal Cortex. Neuron (2017).

8. Yartsev, M. M. & Ulanovsky, N., Representation of three-dimensional space in the hippocampus of flying bats. Science 340, 367–72 (2013).

9. Burak, Y. & Fiete, I. R., Accurate Path Integration in Continuous Attractor Network Models of Grid Cells. PLoS Computational Biology 5, e1000291 (2009).

10. Burgess, N., Barry, C. & O’Keefe, J., An oscillatory interference model of grid cell firing. Hippocampus 17, 801–812 (2007).

11. Fuhs, M. C. & Touretzky, D. S., A Spin Glass Model of Path Integration in Rat Medial Entorhinal Cortex. The Journal of Neuroscience 26, 4266–4276 (2006).

12. Grossberg, S. & Pilly, P. K., How Entorhinal Grid Cells May Learn Multiple Spatial Scales from a Dorsoventral Gradient of Cell Response Rates in a Self-organizing Map. PLoS Computational Biology 8, e1002648 (2012).

13. Navratilova, Z., Giocomo, L. M., Fellous, J., Hasselmo, M. E. & McNaughton, B. L., Phase precession and variable spatial scaling in a periodic attractor map model of medial entorhinal grid cells with realistic after-spike dynamics. Hippocampus 22, 772–789 (2012).

14. Moser, E. I. et al., Grid cells and cortical representation. Nature Reviews Neuroscience 15, 466–481 (2014).

15. Burak, Y. & Fiete, I., Do We Understand the Emergent Dynamics of Grid Cell Activity? The Journal of Neuroscience 26, 9352–9354 (2006).

16. Karlsson, M. P. & Frank, L. M., Awake replay of remote experiences in the hippocampus. Nature Neuroscience 12 (2009).

17. Kudrimoti, H. S., Barnes, C. A. & McNaughton, B. L., Reactivation of hippocampal cell assemblies: effects of behavioral state, experience, and EEG dynamics. The Journal of Neuroscience 19, 4090–101 (1999).

18. O’Neill, J., Senior, T. J., Allen, K., Huxter, J. R. & Csicsvari, J., Reactivation of experience-dependent cell assembly patterns in the hippocampus. Nature Neuroscience 11, 209–215 (2008).

19. Wilson, M. A. & McNaughton, B. L., Reactivation of hippocampal ensemble memories during sleep. Science 265, 676–679 (1994).

20. Brandon, M. P., Bogaard, A. R., Andrews, C. M. & Hasselmo, M. E., Head direction cells in the postsubiculum do not show replay of prior waking sequences during sleep. Hippocampus 22, 604–618 (2012).

21. Peyrache, A., Lacroix, M. M., Petersen, P. C. & Buzsáki, G., Internally organized mechanisms of the head direction sense. Nature Neuroscience 18 (2015).

22. Canto, C. B., Wouterlood, F. G. & Witter, M. P., What Does the Anatomical Organization of the Entorhinal Cortex Tell Us? Neural Plasticity 2008, 381–243 (2008).

23. Fyhn, M., Hafting, T., Treves, A., Moser, M.-B. & Moser, E. I., Hippocampal remapping and grid realignment in entorhinal cortex. Nature 446, 190–4 (2007).

24. Steward, O. & Scoville, S. A., Cells of origin of entorhinal cortical afferents to the hippocampus and fascia dentata of the rat. Journal of Comparative Neurology 169, 347–370 (1976).

25. Zhang, S.-J. et al., Optogenetic dissection of entorhinal-hippocampal functional connectivity. Science 340, 123–2627 (2013).

26. O’Neill, J., Boccara, C. N., Stella, F., Schoenenberger, P. & Csicsvari, J., Superficial layers of the medial entorhinal cortex replay independently of the hippocampus. Science 355, 184–188 (2017).

27. McNaughton, B. L., Battaglia, F. P., Jensen, O., Moser, E. I. & Moser, M.-B., Path integration and the neural basis of the cognitive map. Nature Reviews Neuroscience 7, 663–678 (2006).

28. Welinder, P. E., Burak, Y. & Fiete, I. R., Grid cells: The position code, neural network models of activity, and the problem of learning. Hippocampus 18, 1283–1300 (2008).

29. Kraus, B. J. et al., During Running in Place, Grid Cells Integrate Elapsed Time and Distance Run. Neuron 88, 578-589 (2015).

30. Killian, N. J., Jutras, M. J. & Buffalo, E. A., A map of visual space in the primate entorhinal cortex. Nature 491, 761–4 (2012).

31. Nádasdy, Z., Hirase, H., Czurkó, A., Csicsvari, J. & Buzsáki, G., Replay and time compression of recurring spike sequences in the hippocampus. The Journal of Neuroscience 19, 9497–507 (1999).

32. Lee, A. K. & Wilson, M. A., Memory of Sequential Experience in the Hippocampus during Slow Wave Sleep. Neuron 36, 1183–1194 (2002).

33. Bonnevie, Tora, et al. Grid cells require excitatory drive from the hippocampus. Nature Neuroscience 16, 309–317 (2013).

34. Yoon, K. et al., Specific evidence of low-dimensional continuous attractor dynamics in grid cells. Nature Neuroscience 16, 1077–1084 (2013).

35. Hafting, T., Fyhn, M., Bonnevie, T., Moser, M.-B. & Moser, E. I., Hippocampus-independent phase precession in entorhinal grid cells. Nature 453, 1248-52 (2008).

36. Couey, J. J. et al., Recurrent inhibitory circuitry as a mechanism for grid formation. Nature Neuroscience 16, 318–324 (2013).

37. Pastoll, H., Solanka, L., Rossum, M. & Nolan, M. F., Feedback Inhibition Enables Theta-Nested Gamma Oscillations and Grid Firing Fields. Neuron 77, 141–154 (2013).

38. Dhillon, A. & Jones, R. S. G., Laminar differences in recurrent excitatory transmission in the rat entorhinal cortex in vitro. Neuroscience 99, 413–422 (2000).

39. Fuchs, E. et al., Local and Distant Input Controlling Excitation in Layer II of the Medial Entorhinal Cortex. Neuron 89, 194–208 (2016).

40. Louie, K. & Wilson, M. A., Temporally Structured Replay of Awake Hippocampal Ensemble Activity during Rapid Eye Movement Sleep. Neuron 29, 145–156 (2001).

41. Colgin, L. L., Moser, E. I. & Moser, M.-B., Understanding memory through hippocampal remapping. Trends in Neurosciences 31, 469–77 (2008).

42. Yoon, K., Lewallen, S., Kinkhabwala, A. A., Tank, D. W. & Fiete, I. R., Grid Cell Responses in 1D Environments Assessed as Slices through a 2D Lattice. Neuron 89, 1086–99 (2016).

43. Aronov, D., Nevers, R. & Tank, D. W., Mapping of a non-spatial dimension by the hippocampal–entorhinal circuit. Nature 543, 719–722 (2017).

44. Amari, S.-i., Dynamics of pattern formation in lateral-inhibition type neural fields. Biological Cybernetics 27, 77–87 (1977).

45. Guanella, A., & Verschure, P. F. A model of grid cells based on a path integration mechanism. International Conference on Artificial Neural Networks, 740–749 (2006).

46. Widloski, J. & Fiete, I. R., Cortical microcircuit determination through global perturbation and sparse sampling in grid cells. Preprint at https://www.biorxiv.org/content/early/2015/05/18/019224(2015).

47. Ji, D. & Wilson, M. A., Coordinated memory replay in the visual cortex and hippocampus during sleep. Nature Neuroscience 10, 100-107 (2007).

48. Rothschild, G., Eban, E. & Frank, L. M., A cortical-hippocampal-cortical loop of information processing during memory consolidation. Nature Neuroscience 20, 251–259 (2016).

49. Euston, D. R., Tatsuno, M. & McNaughton, B. L., Fast-Forward Playback of Recent Memory Sequences in Prefrontal Cortex During Sleep. Science 318, 1147–1150 (2007).

50. Ó lafsdóttir, H. F., Carpenter, F. & Barry, C., Coordinated grid and place cell replay during rest. Nature Neuroscience 19, 792–794 (2016).

51. Gothard, K. M., Skaggs, W. E., Moore, K. M. & McNaughton, B. L., Binding of hippocampal CA1 neural activity to multiple reference frames in a landmark-based navigation task. The Journal of Neuroscience 16, 823–35 (1996).

52. Mitchell, S. J. & Ranck, J. B., Generation of theta rhythm in medial entorhinal cortex of freely moving rats. Brain Research 189, 49–66 (1980).

53. Buzsáki, G., Hippocampal sharp waves: Their origin and significance. Brain Research 398, 242–252 (1986).

54. Trimper, J. B., Trettel, S. G., Hwaun, E. & Colgin, L. L., Methodological Caveats in the Detection of Coordinated Replay between Place Cells and Grid Cells. Frontiers in Systems Neuroscience 11, 57 (2017).

55. Zheng, C., Bieri, K., Trettel, S. & Colgin, L., The relationship between gamma frequency and running speed differs for slow and fast gamma rhythms in freely behaving rats. Hippocampus 25, 924–938 (2015).

56. Csicsvari, J., Hirase, H., Czurkó, A., Mamiya, A. & Buzsáki, G., Oscillatory coupling of hippocampal pyramidal cells and interneurons in the behaving Rat. The Journal of Neuroscience 19, 274–87 (1999).

57. Colgin, L. L. et al., Frequency of gamma oscillations routes flow of information in the hippocampus. Nature 462, 353–7 (2009).

58. Tallon-Baudry, C., Bertrand, O., Delpuech, C. & Pernier, J., Oscillatory ?-band (30–70 Hz) activity induced by a visual search task in humans. The Journal of Neuroscience 17 (1997).

59. Geisler, C., Robbe, D., Zugaro, M., Sirota, A. & Buzsáki, G., Hippocampal place cell assemblies are speed-controlled oscillators. Proceedings of the National Academy of Sciences of the United States of America 104, 8149–54 (2007).

